# Allele mining of eukaryotic translation initiation factor genes in *Prunus* species for the identification of new sources of resistance to sharka

**DOI:** 10.1101/2023.01.08.523144

**Authors:** David Tricon, Julie Faivre d’Arcier, Jean-Philippe Eyquard, Shuo Liu, Stéphane Decroocq, Aurélie Chague, Weisheng Liu, Gulnara Balakishiyeva, Alamdar Mamadov, Timur Turdiev, Tatiana Kostritsyna, Bayram M. Asma, Zeynal Akparov, Véronique Decroocq

## Abstract

**Background:** Members of the eukaryotic translation initiation complex are co-opted in viral infection, leading to susceptibility in many crop species, including stone fruit trees (*Prunus spp)*. Therefore, modification of one of those eukaryotic translation initiation factors or changes in their gene expression may result in resistance.

**Objective:** We searched the crop and wild *Prunus* germplasm from the Armeniaca and Amygdalus taxonomic sections for allelic variants in the *eIF4E* and *eIFiso4E* genes, to identify alleles potentially linked to resistance to the *Plum Pox Virus* (PPV).

**Methodology and results:** Over one thousand stone fruit accessions (1,397) were screened for variation in eIF4E and eIFiso4E transcript sequences which are in single copy within the diploid *Prunus* genome. We identified new alleles for both genes that are not evident in haplotypes associated with PPV susceptible individuals. Overall, analyses showed that *eIFiso4E* is genetically more constrained since it displayed less polymorphism than *eIF4E*. We also demonstrated more variation at both loci in the related wild species than in crop species. As the eIFiso4E translation initiation factor was identified as indispensable for PPV infection in *Prunus* species, a selection of ten different *eIFiso4E* haplotypes along 13 individuals were tested by infection with PPV and eight of them displayed a range of reduced susceptibility to resistance, indicating new potential sources of resistance to sharka.

## Introduction

Viral diseases represent an increasing problem in modern, intensive agriculture, and the situation is expected to intensify with global warming. Control measures against viral diseases include the use of virus-free seeds or rootstocks, chemical controls of virus-transmitting vectors, and deployment of virus-resistant cultivars based on dominant or recessive resistance mechanisms (Gómez et al. 2009). While dominant resistance is, in general, an induced and race-specific resistance, recessive resistance in plant-virus interactions is more likely to derive from a passive mechanism due to the absence or to the inappropriate nature of a host factor specifically required by the virus to complete its life cycle (Fraser 1990). The corresponding dominant allele, also called susceptibility allele or (*S)*-gene, is conceptually envisioned as encoding a susceptibility factor needed by the virus (see review Truniger and Aranda 2009). Up to now, resistant cultivars were based, in many cases, on dominant resistance (*R*)-genes which are an attractive option for breeders because they are easy to manipulate in breeding programs. However, they are not always available in the natural diversity of crop species and their interspecific transfer from model plants to crop species proves difficult (Harris et al. 2013; Narusak et al. 2013). An alternative strategy is to employ recessive resistance based on the defect of a (*S)*-gene. Recent studies showed that this resistance mechanism is more easily transferred from model plants to crop species. Indeed, (*S)*-genes are constitutive host cell factors that are co-opted and required by the pathogen to complete or sustain its infectious cycle (*ex*. the translation initiation factors in Le Gall et al. 2011). They are thus expected to be highly conserved across plant genera and if a virus recruits them in a model plant, it likely uses them in its natural host crop species. This is the case of the eukaryotic translation initiation factors, eIF4E and its isoform, eIFiso4E (see review Wang and Krishnaswamy 2012). In consequence, the search for allelic variants of these genes that no longer exhibit a susceptible response, i.e., the type of host variant encountered in a compatible host/virus interaction, could potentially lead to new sources of resistance. This was demonstrated in various crop species such as tomato, melon and pepper. Allele mining by targeting *(S)*-genes in those crop species and natural populations has emerged as an important approach for cloning and characterizing new forms of disease resistance factors (Nieto et al. 2007; Charron al. 2008; Ibiza et al. 2010; Jeong et al. 2012; Gauffier et al. 2016; Poulicard et al. 2016).

In stone fruit tree species, sharka is the most detrimental disease, with significant socio-economic impact, especially in Europe (Cambra et al., 2006). The causative agent is a potyvirus, the *Plum Pox Virus* (PPV) (Rimbaud et al., 2015). Few sources of resistance to this disease have been described (Audergon et al. 1994; Dosba et al. 1994; Escalettes et al. 1998; Martinez-Gomez et al. 2000) but none of them in peach or diploid plum. Resistance to sharka was identified and documented in *Prunus armeniaca* (apricot) (Decroocq et al., 2016) as well as in peach related species, such as *P. davidiana* (Pascal et al. 1998) and *P. dulcis* (almond) (Pascal et al. 2002; Rubio et al. 2003). Although significant effort was devoted to finding genes controlling resistance to PPV, their characterization and utilization have proven to be a long and arduous endeavor (Decroocq et al. 2014). Moreover, those sources of resistance are rather limited, with one single origin per species (Zhebentayeva et al. 2008). Recent history has demonstrated the dangers of relying too heavily on such a limited resistant germplasm, especially when confronted with the diversity of the virus (Rimbaud et al., 2015). To diversify those sources of resistance to PPV and find new ones, other resistance mechanisms were identified in the model plant *Arabidopsis thaliana* that are linked/bind to factors of the translation initiation machinery, eIFiso4E and eIFiso4G1 (Decroocq et al. 2006; Nicaise et al. 2007).

In eukaryotes, translation initiation factors are encoded by a small multi-gene family in which isoforms partly act redundantly. In plants, potyviruses have a specific requirement for a given protein, eIF4E/eIF4G or their isoforms eiFiso4E/eIFiso4G that depends on the host plant and on the virus (Robaglia and Caranta, 2006). For example, *Lettuce mosaic virus* (LMV) uses eIF4E to infect *Lactuca sativa* (lettuce) but uses eIFiso4E in the case of *Arabidopsis thaliana* (Nicaise et al. 2003). Previous studies showed that, in the case of PPV, the eIFiso4E factor is indispensable to viral infection both in *Arabidopsis thaliana* and the European hexaploid plum *P. domestica* (Decroocq et al. 2006; Wang et al. 2013). However, in the diploid plum *P. salicina*, an RNAi silenced *eIFiso4E* transgenic plant could not be obtained. As *eIFiso4E* is a single-copy gene on the *Prunus* diploid genome, this is probably due to a lethal counter-effect of the eIFiso4E null allele on plant growth in diploid *Prunus* species. On the contrary, the silencing of one of the two copies of *eIFiso4G* displays durable and stable resistance to PPV, with no consequence onto plum tree growth (Rubio et al., 2019).

Here, our goal was to investigate natural allelic variation in the *Prunus eIF4E* and *eIFiso4E* genes among the stone fruit cultivated germplasm (apricot, almond and peach crop species) as well as their wild related, ornamental and undomesticated species. Our first objective was to evaluate and compare *eIF4E* and *eIFiso4E* genetic diversity within the wild and the cultivated *Prunus* germplasm, according to their species and to their regions of origin. Several haplotypes with various sites comprising amino acid substitutions, insertions and deletions with however no frame-shift or stop-codon, were identified in both *eIF4E* and *eIFiso4E* sequences. Because most of the crop species of *Prunus* are susceptible to sharka, we examined intraspecific relative to interspecific gene variability to test the hypothesis that more new alleles could be found in wild related, ornamental and undomesticated species. We secondly identified haplotypes of the *eIFiso4E* with variations in the coding sequence that could confer resistance to sharka. We assessed susceptibility to PPV for rare allelic variation of the eIFiso4E susceptible factor with a lower occurrence of presence of 5% and identified new potential sources of resistance to sharka.

## Materials and methods

### Plant material and sampling

The study includes a total of 1,397 individuals from the *Prunus* genus (Table S1). Samples were allocated to three main groups following the subgenera in the *Prunus* taxomony (Figure 1-A). The first group corresponds to representatives of the Armeniaca group (n_Armeniaca_ = 892) divided into three sub-groups representing (i) apricot crop species (*Prunus armeniaca*) (n_Apricot_crop_species_ = 475), (ii) wild, undomesticated apricots (*P. armeniaca*) (n_Wild_apricots_ = 315) and (iii) apricot wild related and ornamental species (n_Apricots_related_species_ = 102). The second main group was Amygdalus (n_Amygdalus_ = 210) divided between (i) almond crop species (*P. dulcis*) (n_Almond_crop_species_ = 138) and (ii) almond wild related species (n_Almonds_related_species_= 72). The last main group was Persica (n_Persica_ = 295) composed of (i) peach crop species (*P. persica*) (n_Peach_crop_species_ = 260) and (ii) peach related species (n_Peach_related_species_= 35).

**Figure 1:**
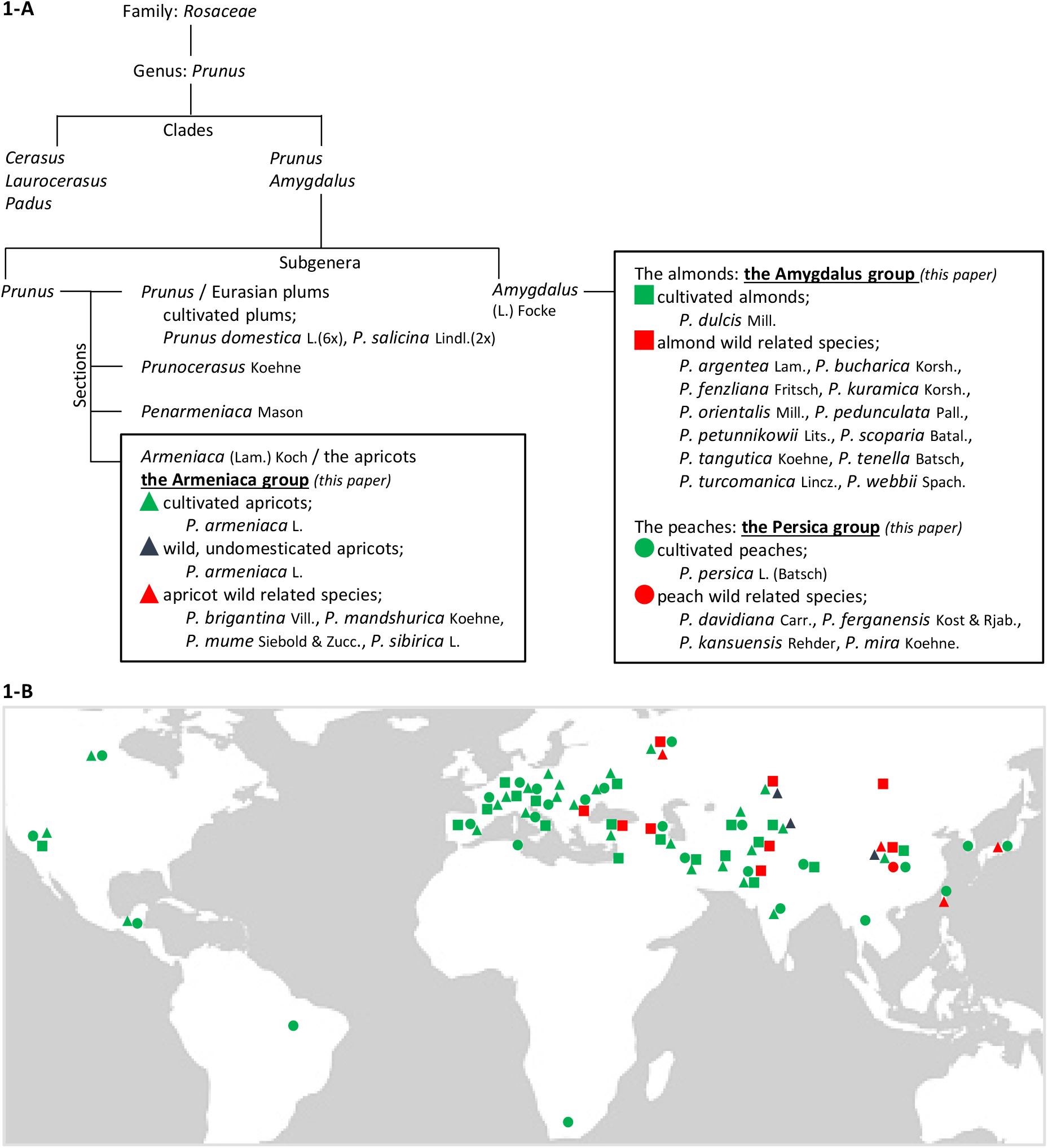
Taxonomy and geographic distribution of *Prunus* accessions **1-A** Taxonomy and schematic phylogeny used in this study. Classification follows Bortiri et al., 2001 linked with Redher (1940) and Mason (1913) previous studies. Black squares localize the three main groups of this work in the *Prunus* genus: the Armeniaca, the Amygdalus and the Persica groups representing accessions related to apricots, almonds and peaches identified by triangles, squares and circles respectively. Green, red and blue colors refer to crop species, wild related species and wild/undomesticated apricots respectively. **1-B** Geographic distribution of the accessions used in this study according to their country of initial sampling (see Table S1).

Apricots, almonds and peaches are crop species that comprise modern varieties, breeding genitors, ancient local varieties or landraces. Wild, undomesticated apricots correspond to individuals sampled away from the cultivated areas in the natural forest mountains of Central Asia. Wild related and ornamental species differ phylogenetically from the *P. armeniaca, P. dulcis* and *P. persica* crop species and were considered as related species of apricots, almonds and peaches, respectively. Individuals were sampled in different geographic areas; in orchards and different germplasm repositories, in private gardens, along the roads, or in natural forest mountains (Figure 1-B). For full details, see Table S1.

### RNA extraction

Around 0.1 to 0.3 mg of fresh or lyophilized young leaves were collected in a 2 ml tube with two iron balls and then stored at -80°C. Samples were ground to obtain a uniform powder. Total RNAs were immediately extracted with the Macherey-Nagel NucleoSpin RNA plant extraction kit (www.mn-net.com). To increase the yield of total RNAs, the extraction procedure was modified by the addition of 1% of beta-mercaptoethanol and 1% w/v of PVP40K in the RAP extraction buffer. Only total RNAs with an absorbance ratio (A_260/280 nm_) of 2 to 2.2 were used to pursue the analysis, otherwise RNAs were re-extracted. RNAs were then stored at -80°C.

### Reverse transcription and genes amplification by PCR

cDNAs were synthesized from total RNAs in 96-well plates with the RevertAid H Minus Reverse Transcriptase and the oligo(dT)_18_ primer according to the manufacturer’s protocol (www.thermoscientific.com).

Specific primers for the amplification of the full-length opening reading frames of the candidate genes were designed according to the published peach genome sequence in the Genomic Database for Rosaceae (GDR, www.rosaceae.org). The candidate genes are referenced as Prupe.4G072600.1 for *eIF4E* (coding sequence length: 705 bp), and Prupe.1G046600.1 for *eIFiso4E* gene (645 bp). For both genes and each individual, polymerase chain reactions (PCRs) were performed in a final volume of 25 µL using the Taq DNA Polymerase from Qiagen (www.qiagen.com) with a modified mix proportions as followed; 10X Buffer/15 mM MgCl2 (1X final), 25 mM MgCl2 (to reach 2.5 mM final), dNTPs mix (0.25 mM final each), forward and reverse primers (0.5 µM final each), 5 U/µl Taq Qiagen (0.625 U per reaction) and 2 µl of cDNAs matrix (20-50 ng/µl). PCR cycling conditions were 3 min at 94°C for general denaturation followed by a one 3-step cycle repeated 40 times (denaturation for 30 sec at 94°C, gene-specific primers’ annealing temperature for 30 sec and extension 40 sec at 72°C) and a final extension for 10 min at 72°C. To perform PCR for *eIF4E*, primers’ annealing temperature was optimized at 61°C with the forward primer 5’-CGCCAAGAAAGAAAAGCGAG-3’ and the reverse primer 5’-GCAAAGAACAATATACACATCA-3’ and for *eIFiso4E* the annealing step was performed at 58°C with the forward primer 5’-AAACAACACAACCCCGACAG-3’ and the reverse primer 5’-TCAAACATTGTATCGA-3’. PCR products were verified by electrophoresis with a 1.5% agarose gel and visually quantified by the MassRuler DNA ladder from ThermoScientific.

### Allelic sequencing, phasing and alignment

PCR products were sequenced with the Sanger method by the Genewiz company, following recommendations available on the www.genewiz.com website. Quality of sequences was verified from the chromatograms using Chromas v.2.5.1 (https://technelysium.com.au/wp/chromas/). Sequences of individuals were all classified in a data file corresponding to the appropriate main group and sub-group in which they belonged (see Plant material and sampling section). As the *Prunus* species used in the current study are all diploid, each gene can have up to two alleles. In this case, heterozygous allelic forms with unphased genotypic data were rebuilt with the ELB algorithm method implemented in Arlequin v.3.5.2.2 as a pseudo-Bayesian approach to specifically estimate gametic phase in recombining sequences (Excoffier et al., 2010). All phased haplotypes were first aligned altogether using ClustalW Multiple Alignment method (1,000 Bootstraps) using BioEdit v.7.1.3.0 software (Hall 1999), then trimmed to deal with nothing else than the coding sequence from the start to the stop codons and finally translated into amino acid sequences.

### Polymorphism detection and statistical analyses

For each main group and sub-group, amino acid sequences were compared to a reference haplotype corresponding to the sequence coming from a susceptible individual to PPV i.e., the apricot cultivar ‘Moniqui’ (*P. armeniaca*), the almond cultivar ‘Aï’ (*P. dulcis*) and the peach rootstock ‘GF305’ (*P. persica*) for the Armeniaca, Amygdalus and Persica groups and sub-groups respectively (see Plant material and sampling section). These sequences of reference were respectively called Arm_P00, Amy_P00 and Per_P00 (“P” for protein) with the prefix “eIF4E_” or “eIFiso4E_” (a.k.a. eIF4E_Arm_P00). For each new haplotype detected, the number after “P” was incremented (“P01”, “P02”, …).

For both coding and amino acid sequences levels of homozygosity, heterozygosity, haplotypic richness and the presence of variations were evaluated and compared between groups and sub-groups using Chi-square tests (*χ*^*2*^ tests) statistical analyses performed by XLStat software v.2020.1.3. Frequencies of haplotypes and amino acid variations were also calculated and were called rare haplotypes and rare amino acid variations when their occurrences were less than 5%.

### Genetic diversity parameters

Genetic diversity estimates were calculated using DNAsp v.5.10.01 software (Librado and Rozas 2009). The haplotype diversity (*Hd*), the nucleotide diversity (*π*), the ratio of non-synonymous to synonymous substitutions (*dn/ds* ratio) and the Tajima’s *D* statistic were calculated from allelic data for each main group and sub-group. The Datamonkey online database (www.datamonkey.org) was used to test the selection pressure on (i) the whole genes sequences with the RELAX test to analyze whether the strength of the selection has been relaxed or intensified along the phylogeny of the sequences and the BUSTED model (Branch-site Unrestricted Statistical Test for Episodic Diversification) to provide a gene-wide test for positive selection at at least one site on at least one phylogenetic branch, and (ii) at individual sites with the FUBAR test (Fast Unconstrained Bayesian AppRoximation) for large data sets with a Bayesian approach to infer the *dn/ds* ratio on a per-site basis and detect positive or negative pervasive selection at the amino acid level assuming the selection pressure for each site is constant along the entire phylogeny (Weaver et al., 2018).

### Predicting effects of amino acid variations on protein conformation/function

To predict the potential effect of amino acid variations on the protein conformation and/or function, three computational methods were used: (i) Meta-SNP (https://snps.biofold.org/meta-snp/index.html) a random forest-based binary classifier predictor combining predictions of four methods (SNAP2, SIFT, PANTHER, PhD-SNP) and four elements extracted from the PhD-SNP protein sequence profile based on training dataset derived from SwissVar (Capriotti et al., 2013), (ii) MutPred2 (http://mutpred.mutdb.org) an algorithm able to quantify the pathogenicity of amino acid substitutions and describe how they can affect the protein function by modeling a broad repertoire of structural and functional alterations from amino acid sequence (Pejaver et al., 2020) and (iii) PredictSNP (https://loschmidt.chemi.muni.cz/predictsnp1/) a consensus classifier combining eight prediction methods (MAPP, PhD-SNP, PolyPhen-1/-2, SIFT, SNAP, nsSNPAnalyser, PANTHER) to provide a more accurate and robust alternative to the predictions based on individual integrated tools and weighted by the method-specific confidence scores (Bendl et al., 2014). Even though these softwares based their predictions on mammal (mostly humans) databases (no such plant-specific predictors exist), they enabled the classification of amino acid variations found along the eIFiso4E protein for Armeniaca, Amygdalus and Persica individuals to select variable haplotypes for testing PPV infection in the greenhouse.

### Phenotypic evaluation

Once the selection of individuals with variable haplotypes was established, phenotypic evaluation of PPV resistance was performed in a high confinement greenhouse following the protocol of Decroocq et al 2016. Three replicates per accession were tested over three vegetative cycles of observations with two rounds of measurements each by serological assays (ELISA). Indeed, to look for the presence of the virus on the leaves, optical density values (OD) were measured each time by ELISA. When OD was at least twice higher than the OD value of the negative control (the non-infected cultivar GF305), the sample was considered as infected. In this case, a score of 1 was attributed to the sample. Otherwise, the score was 0. An average response score was calculated after each cycle and a global average score was obtained at the end of the three cycles. Accessions were considered resistant with a global score of 0. Three levels of susceptibility were then categorized with a weak (0 < score < 0.200), a moderate (0.200 ≤ score < 0.500) and a high (0.500 ≤ score ≤ 1) score.

## Results

### *eIF4E* and *eIFiso4E* coding and amino acid sequence lengths

The *eIF4E* coding sequence was 702 base pairs long, from the start codon to the stop codon and the predicted protein was 234 amino acids long for the three groups. Sequence length variations were observed for *eIFiso4E* due to the presence or the absence of triplets of nucleotides causing no frame shift. In consequence, the *eIFiso4E* coding sequence varies from 639 to 648 base pairs and the protein, from 213 to 216 amino acids (Table 1).

**Table 1:** *eIF4E* and *eIFiso4E* number of individuals used for data analyses, lengths of coding (in base pairs [bp]) and amino acid sequences identified in the main groups and sub-groups

|  | Number of individuals |  | Coding sequence length (bp) |  | Amino acids length |  |
| --- | --- | --- | --- | --- | --- | --- |
|  | *eIF4E* | *eIFiso4E* | *eIF4E* | *eIFiso4E* | *eIF4E* | *eIFiso4E* |
| Armeniaca group | 695 | 767 | 702 | 639-642-645 | 234 | 213-214-215 |
| Apricot crop species | 400 | 413 | 702 | 639-642-645 | 234 | 213-214-215 |
| Wild apricots | 220 | 268 | 702 | 639-642 | 234 | 213-214 |
| Apricot related species | 75 | 86 | 702 | 642 | 234 | 214 |
| Amygdalus group | 189 | 182 | 702 | 642-645-648 | 234 | 214-215-216 |
| Almond crop species | 134 | 127 | 702 | 642-645-648 | 234 | 214-215-216 |
| Almond related species | 55 | 55 | 702 | 642-645-648 | 234 | 214-215-216 |
| Persica group | 284 | 282 | 702 | 642-645-648 | 234 | 214-215-216 |
| Peach crop species | 252 | 248 | 702 | 642-645-648 | 234 | 214-215-216 |
| Peach related species | 32 | 34 | 702 | 642-645 | 234 | 214-215 |

### Polymorphism and heterozygosity among the *eIF4E* and *eIFiso4E* loci

Overall data and results are illustrated and reported in Figures 2-A & 2-B and Tables S2 & S3.

**Figure 2:**
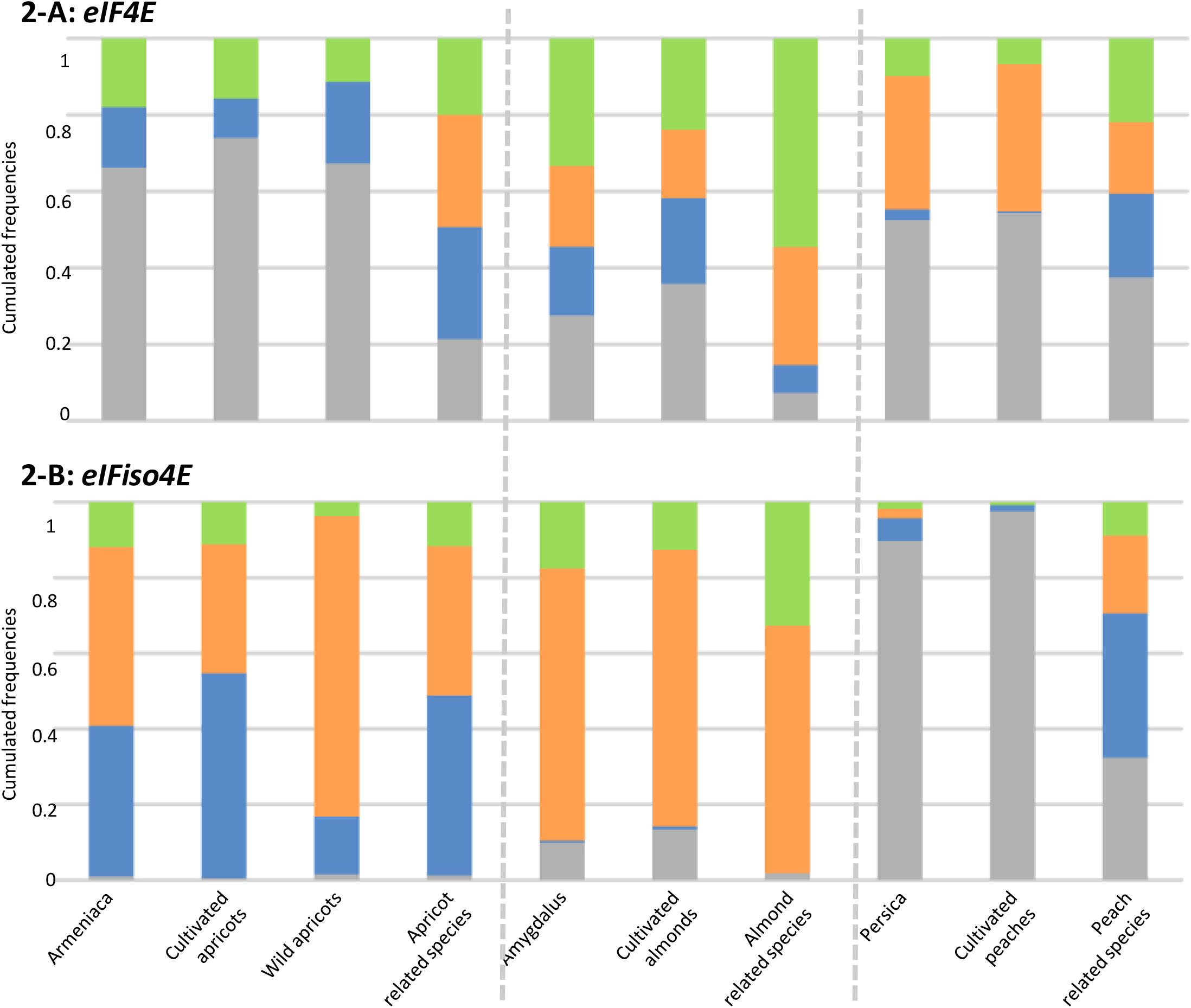
Stacked diagrams of cumulated frequencies following the classification of the accessions in the main groups and in the sub-groups regarding **figure 2-A** for *eIF4E* and **figure 2-B** for *eIFiso4E* amino acid sequences compared to their own reference sequence. Diagrams categorize frequencies following accessions ; with similar sequences to the reference sequence in grey, with at least one nucleotide substitution resulting in a synonymous variation in blue, with at least one amino acid variation (occurrence of presence > 5%) in orange and, with at least one rare amino acid variation (occurrence of presence < 5%) in green.

At the *eIF4E* locus, regardless of the main groups, wild related species showed a significantly higher allelic and haplotypic richness (*p*-values < 0.001) than crop species. Although data were not significantly different in few cases, the wild related species tend to display higher heterozygosity and a higher number of individuals with nucleotide substitutions and amino acid changes than the crop species (Table S2-B). Moreover, all diversity parameters (substitutions/variations and heterozygosity) were the highest in the Amygdalus group compared to the Armeniaca and Persica groups (*p*-values < 0.0001, Table S2-B).

At the *eIFiso4E* locus, wild related species for the three main groups also showed significantly higher allelic and haplotypic richness than the crop species (*p*-values < 0.05, Table S3-B). Moreover, as for *eIF4E*, wild related species showed a significantly higher heterozygosity level than the crop species, except for the Amygdalus group. While the frequency of individuals with variations in coding sequences compared to the reference individual is not significantly different between the three Armeniaca subgroups, in contrast amino acid variations are significantly higher in wild apricots (*p-*value < 0.0001, Table S3-B). The most striking observation was for the Persica group in which wild related species showed a very significantly higher diversity than crop species (*p-*values < 0.0001 for all measured criteria) (Table S3-B).

The three main groups, i.e., Armeniaca, Amygdalus and Persica, showed significant differences for all parameters of diversity of *eIFiso4E* displayed in Table S3-B (*p*-values < 0.0001). The Amygdalus group showed both the highest number of individuals with amino acid variation and haplotypic richness while the Armeniaca group had the highest number of individuals with heterozygosity and variations at the coding sequence level. The Persica group showed the lowest diversity for each criterion.

### Genetic diversity of *eIF4E* and *eIFiso4E* among the three *Prunus* groups

Nucleotide (*π*) and haplotype (*Hd*) diversities were both calculated across the *eIF4E* and *eIFiso4E* gene sequences for the Armeniaca, Amygdalus and Persica groups (Table 2-A).

**Table 2:** *eIF4E* and *eIFiso4E* genetic diversity parameters **2-A** Values of nucleotide diversity (*π*), haplotype diversity (*Hd*), ratio of substitution rate at non-synonymous and synonymous sites (*dn/ds*), and selection Tajima’s *D* statistics with corresponding significance values (* for *p* ≤ 0.05; ** for *p* ≤ 0.01, calculated with DNAsp v.5.10.01 software in the main groups and sub-groups; **2-B** Ratios of *π, Hd*, and *dn/ds* between the main groups and their sub-groups

| Table 2-A | $\pi$ | | $Hd$ | | $dn/ds$ | | Tajima's $D$ | |
| --- | --- | --- | --- | --- | --- | --- | --- | --- |
|  | *eIF4E* | *eIFiso4E* | *eIF4E* | *eIFiso4E* | *eIF4E* | *eIFiso4E* | *eIF4E* | *eIFiso4E* |
| Armeniaca group | 0.00126 | 0.0029 | 0.3789 | 0.8016 | 0.27891 | 0.11826 | -2.28525 | -1.6838 |
| Apricot crop species | 0.00116 | 0.00279 | 0.3198 | 0.8293 | 0.55307 | 0.08791 | -2.28222 ** | -1.24793 |
| Wild apricots | 0.00078 | 0.00201 | 0.3210 | 0.6117 | 0.09455 | 0.18278 | -1.71567 | 0.20376 |
| Apricot related species | 0.00286 | 0.00450 | 0.7417 | 0.8910 | 0.15252 | 0.07720 | -1.57057 | -1.28773 |
| Amygdalus group | 0.00503 | 0.00516 | 0.8160 | 0.9180 | 0.26722 | 0.15331 | -2.05692 * | -1.74178 |
| Almond crop species | 0.00255 | 0.00375 | 0.7308 | 0.8870 | 0.26898 | 0.13495 | -2.10693 * | -1.59196 |
| Almond related species | 0.00839 | 0.00634 | 0.9046 | 0.8537 | 0.32982 | 0.17519 | -1.56449 | -1.29035 |
| Persica group | 0.00142 | 0.00082 | 0.5752 | 0.1889 | 0.63500 | 0.03858 | -1.90355 * | -2.10157 ** |
| Peach crop species | 0.00099 | 0.00022 | 0.5395 | 0.0438 | 2.45652 | 0.03191 | -1.83100 * | -1.97602 * |
| Peach related species | 0.00404 | 0.00313 | 0.7267 | 0.7401 | 0.16279 | 0.07125 | -0.77135 | -1.29487 |
| Table 2-B | *eIF4E* |  | *eIFiso4E* |  |  |  |  |  |
| | ratio of $\pi$ | ratio of $Hd$ | ratio of $\pi$ | ratio of $Hd$ | | | | |
| Amygdalus / Armeniaca | 3.99206 | 2.15360 | 1.77931 | 1.14521 |  |  |  |  |
| Amygdalus / Persica | 3.54225 | 1.41864 | 6.29268 | 4.85971 |  |  |  |  |
| Armeniaca / Persica | 0.88732 | 0.65873 | 3.53659 | 4.24351 |  |  |  |  |
| Apricot related species / Apricot crop species | 2.46552 | 2.31926 | 1.61290 | 1.07440 |  |  |  |  |
| Apricot related species / wild apricots | 3.66667 | 2.31059 | 2.23881 | 1.45660 |  |  |  |  |
| Almond related species / Almond crop species | 3.29020 | 1.23782 | 1.69067 | 0.96246 |  |  |  |  |
| Peach related species / Peach crop species | 4.08081 | 1.34699 | 14.22727 | 16.89726 |  |  |  |  |

For both genes, *π* values were globally in the same range (10^−3^) except for the wild apricots for *eIF4E* and for the peach crop species for both genes (10^−4^). Similar ranges of nucleotide diversity have already been observed in whole genome sequences of apricot and peach crop and wild species (Groppi et al., 2021, Cao et al., 2019). In *eIFiso4E, Hd* was in the same range of values (10^−1^) as *eIF4E* except for the peach crop species sub-group (10^−2^). For both genes, *π* and *Hd* values were higher in all wild related species sub-groups compared to the crop species ones, except, the *Hd* value for *eIFiso4E* in the almond crop species where it was slightly higher than that of the wild related species. Moreover, the Amygdalus group showed the highest values while the lowest ones were observed in the Persica group except for *Hd* of *eIF4E* in the Armeniaca group. Additionally, peach crop species displayed a significant drop in both *π* and *Hd eIFiso4E* values in comparison with its wild related species (Tables 2-A, 2-B)

The Tajima’s *D* statistics showed a negative value in all *Prunus* groups except for a slight neutral value at the *eIFiso4E* locus in wild apricots (e.g., 0.20376, Table 2-A). The non-synonymous/synonymous substitution ratio, *dn/ds*, was found less than 1, showing evidence that synonymous mutations are more frequent than the non-synonymous ones, excepted in *eIF4E* for peach crop species (*dn/ds* = 2.45652, Table 2-A). These ratio values also indicated that non-synomymous mutations were less prevalent on *eIFiso4E* compared to *eIF4E*, in particular in crop and wild related species of Armeniaca and Persica (*dn/ds*_*_eIFiso4E*_ < *dn/ds*_*_eIF4E*_, Table 2-A). Moreover, at the whole gene level in all groups and sub-groups, for both *eIF4E* and *eIFiso4E* the RELAX and BUSTED tests did not reveal any evidence of gene-wide relaxed or intensified selection, neither positive or negative selection in the phylogeny of the genetic diversity of the sequences (*p-*values > 0.05).

### *eIF4E* and *eIFiso4E* haplotypes

At the amino acid sequence level, a total of 49 *eIFiso4E* haplotypes differing from the PPV susceptible haplotypes of the Armeniaca, Amygdalus and Persica groups were identified, whereas 99 haplotypes were counted for *eIF4E* (Table 3-A). It was shown previously that PPV infection requires a functional eIFiso4F translation initiation complex (Wang et al, 2013; Rubio et al, 2019). In consequence, we will hereafter focus exclusively on eIFiso4E haplotype variation. Susceptibility to the virus is predominant in the *Prunus* germplasm, therefore, susceptibility alleles of *eIFiso4E* are expected to be more common than the rare resistant alleles among the *Prunus* germplasm.

**Table 3:** Number of individuals in the different classes of haplotypes and following the number of amino acid variations along the eIF4E and eIFiso4E sequences **3-A** eIF4E and eIFiso4E class of haplotypes derived from the susceptible reference in the main groups regarding occurrence (< 5%), homozygosity and localization of amino acid variation in the interaction domains (DI or DII) of the sequences. Numbers underlined indicate the number of classes of haplotypes found with an occurrence < 5% among the total number of classes. Numbers in brackets indicate the number of classes of haplotypes found homozygous among the total number of classes. Numbers in red color indicate the number of classes of haplotypes with amino acid variation in the interaction domains found homozygous and with an occurrence < 5%; **3-B** Number of non-synonymous positions and amino acid variations along *eiF4E* and *eIFiso4E* sequences regarding occurrence (< 5%), homozygosity, and localization of the amino acid variation in the interaction domains (DI or DII) of the proteins. Numbers underlined indicate the number of amino acid variations found with an occurrence < 5%. Numbers in brackets indicate the number of amino acid variation found homozygous along the whole sequence. Numbers in red color indicate the number of amino acid variations in the interaction domains found homozygous and with an occurrence < 5%.

| Table 3-A |  |  |  |  |  |  |  |  |
| --- | --- | --- | --- | --- | --- | --- | --- | --- |
| Number of variable haplotypes: | Armeniaca |  | Amygdalus |  | Persica |  | 3 Prunus groups |  |
|  | *eIF4E* | *eIFiso4E* | *eIF4E* | *eIFiso4E* | *eIF4E* | *eIFiso4E* | *eIF4E* | *eIFiso4E* |
| Differing from the susceptible reference haplotype | 41 | 17 | 44 | 24 | 14 | 8 | 99 | 49 |
| With occurrence < 5% | 41 | 16 | 42 | 20 | 13 | 8 | 96 | 44 |
| With occurrence < 5% and homozygous | 12 | 8 | 12 | 10 | 8 | 5 | 32 | 23 |
| With amino acid variation(s) inside the interaction domains | 12 / <u>12</u> (6) | 1 / <u>1</u> (1) | 21 / <u>20</u> (6) | 4 / <u>4</u> (3) | 3 / <u>3</u> (3) | 3 / <u>3</u> (2) |  |  |
| With amino acid variation(s) inside interaction domain I | 2 / <u>2</u> (0) | 1 / <u>1</u> (1) | 0 | 1 / <u>1</u> (0) | 0 | 0 |  |  |
| With amino acid variation(s) inside interaction domain II | 10 / <u>10</u> (6) | 0 | 21 / <u>20</u> (6 / <u>5</u>) | 3 / <u>3</u> (3) | 3 / <u>3</u> (3) | 3 / <u>3</u> (2) |  |  |
| Table 3-B |  |  |  |  |  |  |  |  |
| Number of: | Armeniaca |  | Amygdalus |  | Persica |  |  |  |
|  | *eIF4E* | *eIFiso4E* | *eIF4E* | *eIFiso4E* | *eIF4E* | *eIFiso4E* |  |  |
| Positions with non-synonymous variations along the amino acid sequence | 32 | 19 | 32 | 20 | 16 | 8 |  |  |
| Amino acid variations | 34 | 22 | 40 | 24 | 16 | 9 |  |  |
| Amino acid variations with occurrence < 5% | 34 | 21 | 34 | 19 | 15 | 9 |  |  |
| Homozygous amino acid variations with occurrence < 5% | 13 | 14 | 15 | 10 | 10 | 6 |  |  |
| Amino acid variations inside the interaction domains | 4 / <u>4</u> (1) | 2 / <u>2</u> (2) | 1 / <u>0</u> (1) | 3 / <u>3</u> (2) | 1 / <u>1</u> (1) | 1 / <u>1</u> (1) |  |  |
| Amino acid variations inside interaction domain I | 2 / <u>2</u> (0) | 2 / <u>2</u> (2) | 0 | 1 / <u>1</u> (0) | 0 | 0 |  |  |
| Amino acid variations inside interaction domain II | 2 / <u>2</u> (1) | 0 | 1 / <u>0</u> (1) | 2 / <u>2</u> (2) | 1 / <u>1</u> (1) | 1 / <u>1</u> (1) |  |  |

Details of amino acid variations in haplotypes observed among the *Prunus* germplasm in the Armeniaca, Amygdalus and Persica groups are respectively depicted in Tables S4-A, S4-B, S4-C for *eIFiso4E*. As mentioned above, we did not comment data obtained for *eIF4E* (Tables S5-A, S5-B, S5-C). Indeed, we found that the *eIFiso4E* susceptible reference haplotypes are more prevalent than other haplotype frequencies in Persica (0.968085, Table S4-C) and in Armeniaca (0.565189, Table S4-A) and as well as in Amygdalus (0.456054, Table S4-B), if we overlook length polymorphism with the loss of one Alanine at position 28 (A28-) for *eIFiso4E*_Amy_P01.

Attention was first given to classes of haplotypes found with a frequency lower than 5% in each main group (also called rare haplotypes) as reported in Table 3-A. Rare haplotypes were identified both in the crop and the wild related species sub-groups. EIFiso4E haplotypes differing from the susceptible reference were more often native to Asia (Central and Eastern Asia) and to the irano-caucasian region for Armeniaca (Table S4-A) and Amygdalus with a Western European additional contribution for the latter (Table S4-B). The few diverging Persica haplotypes came from Eastern Asia (Table S4-C).

Several positions along the amino acid sequences were identified with at least one non-synonymous variation compared to the chosen reference sequence; 19, 20 and 8 positions for Armeniaca, Amygdalus and Persica groups respectively (Table 3-B). *EIFiso4E* is a susceptibility gene for which a homozygous mutation can lead to recessive resistance to PPV (Decroocq et al, 2006), thus, we focused on homozygous variations over the *eIFiso4E* coding sequence. Among them, rare amino acid variations (occurrence < 5%) were selected under this threshold since resistance is less frequent than susceptibility in *Prunus* germplasm (Pascal et al, 2002; Marandel et al, 2009) (Tables 3-A, 3-B). We further looked for variations within the two interaction domains, domain I (DI) and domain II (DII), involved in the plant/virus compatible interaction as described by Charon et al. (2008) (Tables S4 & S5 and Figures 3-A, 3-B). Along the eIFiso4E amino acid sequence, DI was delimited from position 46 to position 66 and DII from position 93 to position 96 (Figure 3-A). Additionally, we focused on variations impacting potentially the eIFiso4E 3D conformation and functionality. In the latter case, Meta-SNP, MutPred2 and PredictSNP online predictors were used to predict the impact of the variation(s) at the molecular and functional levels of the protein (Tables S6-A, S6-B, S6-C).

**Figure 3:**
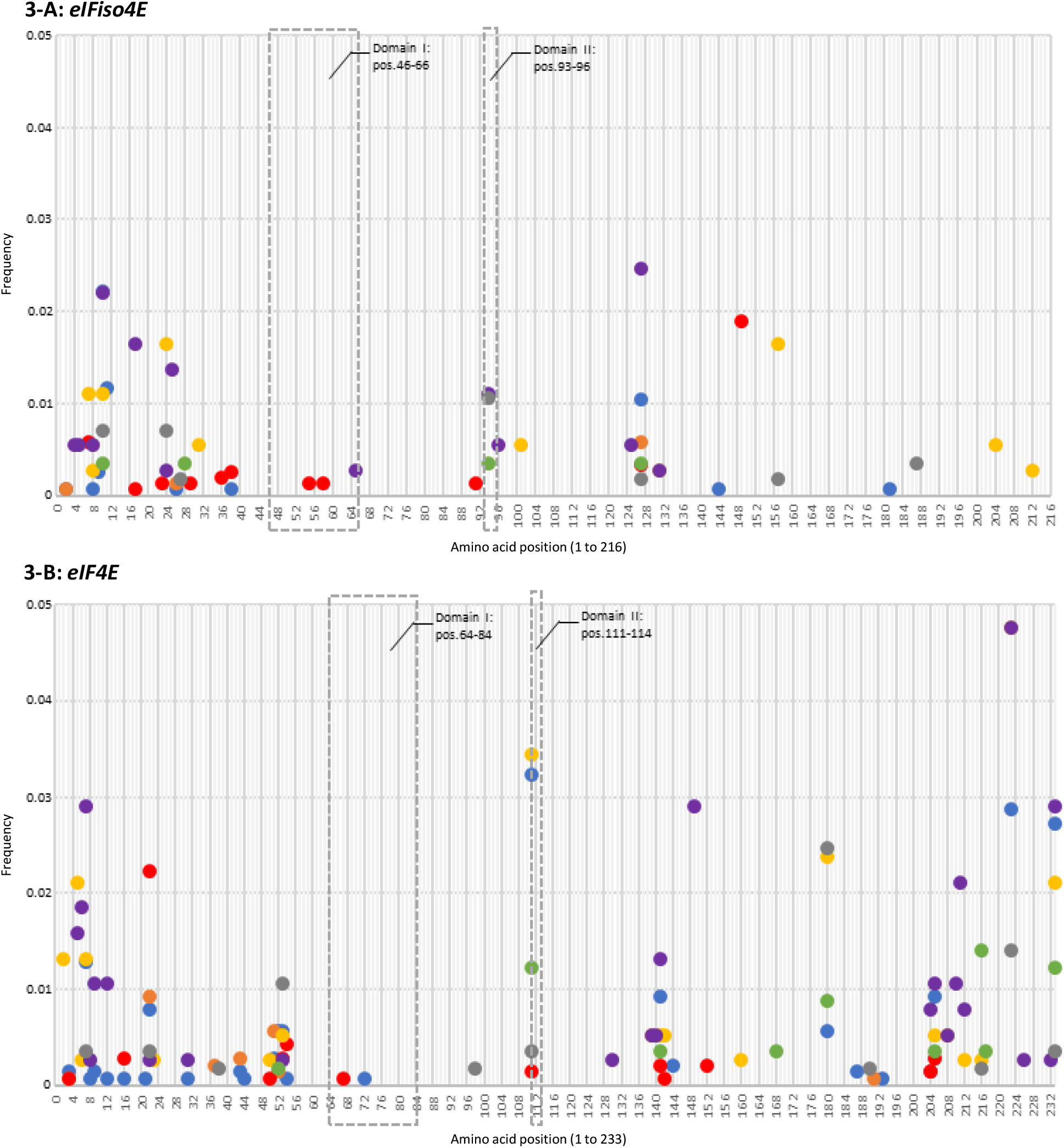
Frequency of rare amino acid variations (occurrence of presence < 5%) along *eIFiso4E* (**Figure 3-A**) and *eIF4E* (**Figure 3-B**) sequences. Interaction domains with the virus: Domain I and Domain II are localized inside dotted grey rectangles. Each coloured circle represents one amino acid variation; each color refers to one sub-group as apricot crop species (blue), wild/undomesticated apricots (orange), apricot wild related species (red), almond crop species (yellow), almond wild related species (purple), peach crop species (green) and peach wild related species (grey).

Regarding all the groups at the same time, one hot spots’ region of rare amino acid variations was identified along the *eIFiso4E* sequences, in the N-terminal region, within the first forty amino acids of the sequence whereby up to 90% of the non-synonymous rare variations mapped in this region (Figure 3-A, Tables S4). Other variations were scattered along the amino acid sequence.

The analysis of the selection pressure for each site performed by the FUBAR test identified one amino acid variation per group that was positively selected (*p-*values < 0.05): D149E for Armeniaca, A127P for Amygdalus and N94S in DII for Persica.

In the Armeniaca group, the eIFiso4E_Arm_P15 haplotype was the only one described with homozygous amino acid variations in the interaction domain I at positions 55 and 58 (K55N and Q58K, respectively) (Table 3-A). It was identified within the US_108 accession (cv. Yanmei, a *P. mume* accession) originating from Japan (Tables 3-A, 3-B, S4-A, S6-A). Together with the K55N amino acid variation, four other variations at positions 2, 144 and 181 (A2E, A2V, Q144H and I181N) were predicted to have an effect in protein conformation. However, they were found in heterozygous allelic forms in the eIFiso4E_Arm_P04, eIFiso4E_Arm_P06, eIFiso4E_Arm_P09 and eIFiso4E_Arm_P17 haplotypes, respectively (Table S6-A).

In the Amygdalus group, three different variations in the interaction domains were identified; S65T (DI) in the heterozygous eIFiso4E_Amy_P16, N94S (DII) in the homozygous eIFiso4E_Amy_P22 and eIFiso4E_Amy_P24, and D96E (DII) in the homozygous eIFiso4E_Amy_P21 with a potential effect on protein conformation for the latter (Tables 3-A, 3B, S4B, S6B). Homozygous variations in DII were found in the almond related species *P. petunnikowii* (US_073 and US_189) for N94S *and P. tenella* (US_135) for D96E (Table S6-B). Those three individuals came from Kazakhstan in central Asia (Table S1). Predictions of non-synonymous variations with a significant effect on protein conformation at other positions as R101G, T131I, T131P, K204I and R212L were all detected in heterozygous allelic forms (Tables S4-B, S6-B).

In the Persica group, four haplotypes were detected with the variation N94S in DII (Table S4-C). This amino acid variation was found homozygous in eIFiso4E_Per_P01 and eIFiso4E_Per_P04 from the *P. persica* AZ_092 (Azerbaïjan) and the *P. davidiana* CH_002 (China) accessions, respectively. It was also found heterozygous in the eIFiso4E_Per_P04 and eIFiso4E_Per_P07 haplotypes (Tables S6-C). One single non-synonymous variation, E187D, that could result in a significant change in protein conformation was identified homozygous in the haplotype eIFiso4E_Per_P06 from the CH_154_3 accession (*P. davidiana*, China) (Tables S1, S6-C).

The variation A127P was observed in all the three *Prunus* groups regarding the reference sequences (Figure 3-A) and was found homozygous in the Amygdalus *P. orientalis* TR_114, TR_115 (eIFiso4E_Amy_P05) and *P. webbii* US_193 accessions (eIFiso4E_Amy_P18). In Armeniaca, it was present in the homozygous eIFiso4E_Arm_P03 with KZ_230_3 while in Persica, it was found heterozygous in the *P. davidiana* FR_AVI_056 accession (eIFiso4E_Per_P08).

Based on the above criteria a.k.a. non-synonymous and homozygous variation(s) potentially affecting the protein conformation of eIFiso4E and prioritizing the interaction domains, we selected a list of accessions to test for resistance to PPV under controlled conditions in a high confinement greenhouse (Table 4). Amino acid variations G38R in Armeniaca, A31P and S157N in Amygdalus were also targeted as they were homozygous and/or rare variations.

**Table 4:**
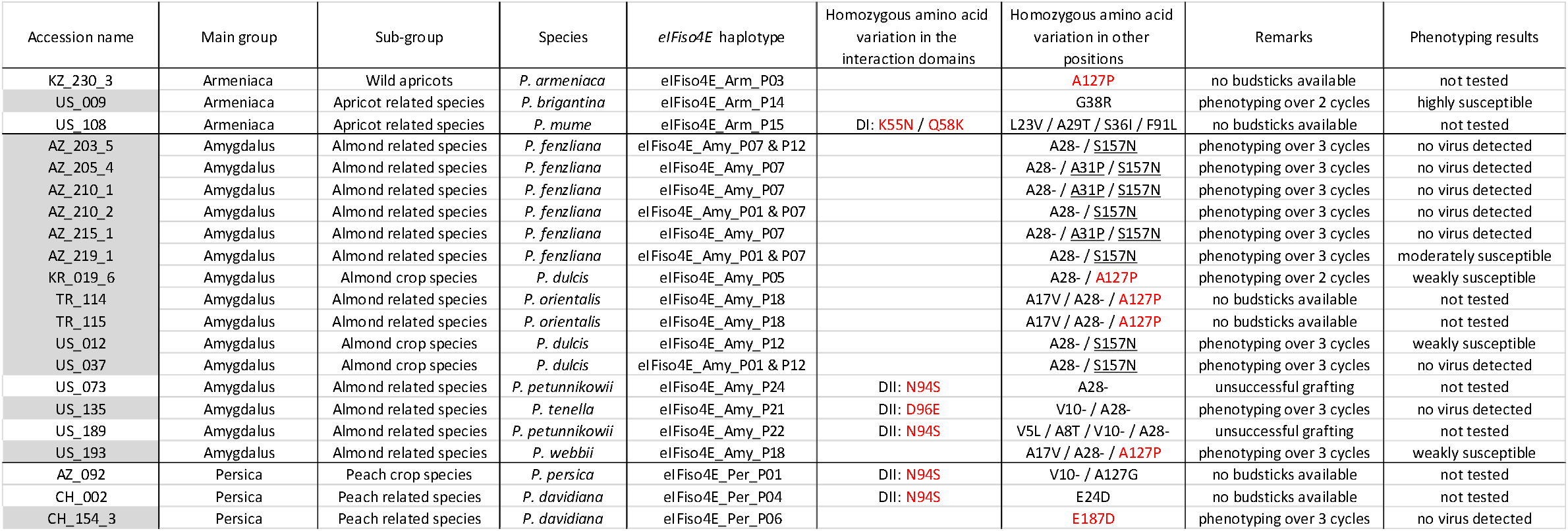
List of selected accessions with eIFiso4E haplotypes derived from their own reference sequence. Selection was done following several criteria (i) a homozygous amino acid variation as a mandatory criterion, (ii) an amino acid variation in at least one interaction domain and/or with a significant predicted effect (see Tables S6), (iii) a rare amino acid variation (occurrence < 5%) and (iv) accessions with budsticks available to be grafted. Accessions highlighted in grey were actually phenotyped for PPV resistance in high confinement greenhouse. One amino acid variation with a significant predicted effect on protein conformation/function is highlighted in red. Other amino acid variations tested are underlined. The symbol “-” represents an amino acid deletion.

In total, thirteen accessions were phenotyped up to three successive vegetative cycles (Table 4). The presence/absence of viral particles was estimated by serological assays (ELISA) using a PPV-specific antibody. No virus was detected over three cycles for seven Amygdalus accessions: one almond crop species *P. dulcis* US_037 and six almond wild related species among which five representatives of *P. fenzliana* (AZ_203_5, AZ_205_4, AZ_210_1, AZ_210_2, AZ_215_1) and one *P. tenella* (US_135). One representative of the peach wild related species, *P. davidiana* CH_154_3 was also scored PPV negative (Figure 4). Intermediate susceptibility levels were measured in Amygdalus *for P. dulcis* KR_019_6 and US_012 as well as in *P. webbii* US_193. Amygdalus *P. fenzliana* AZ_219_1 and Armeniaca *P. brigantina* US_009 were moderately to highly susceptible, respectively (Table 4, Figure 4, Table S7). In total, seven accessions were resistant to sharka. They will be further studied in progenies from crosses of these resistant accessions with PPV susceptible accessions, to verify co-segregation between the newly identified allelic variations in *eIFiso4E* and resistance to sharka.

**Figure 4:**
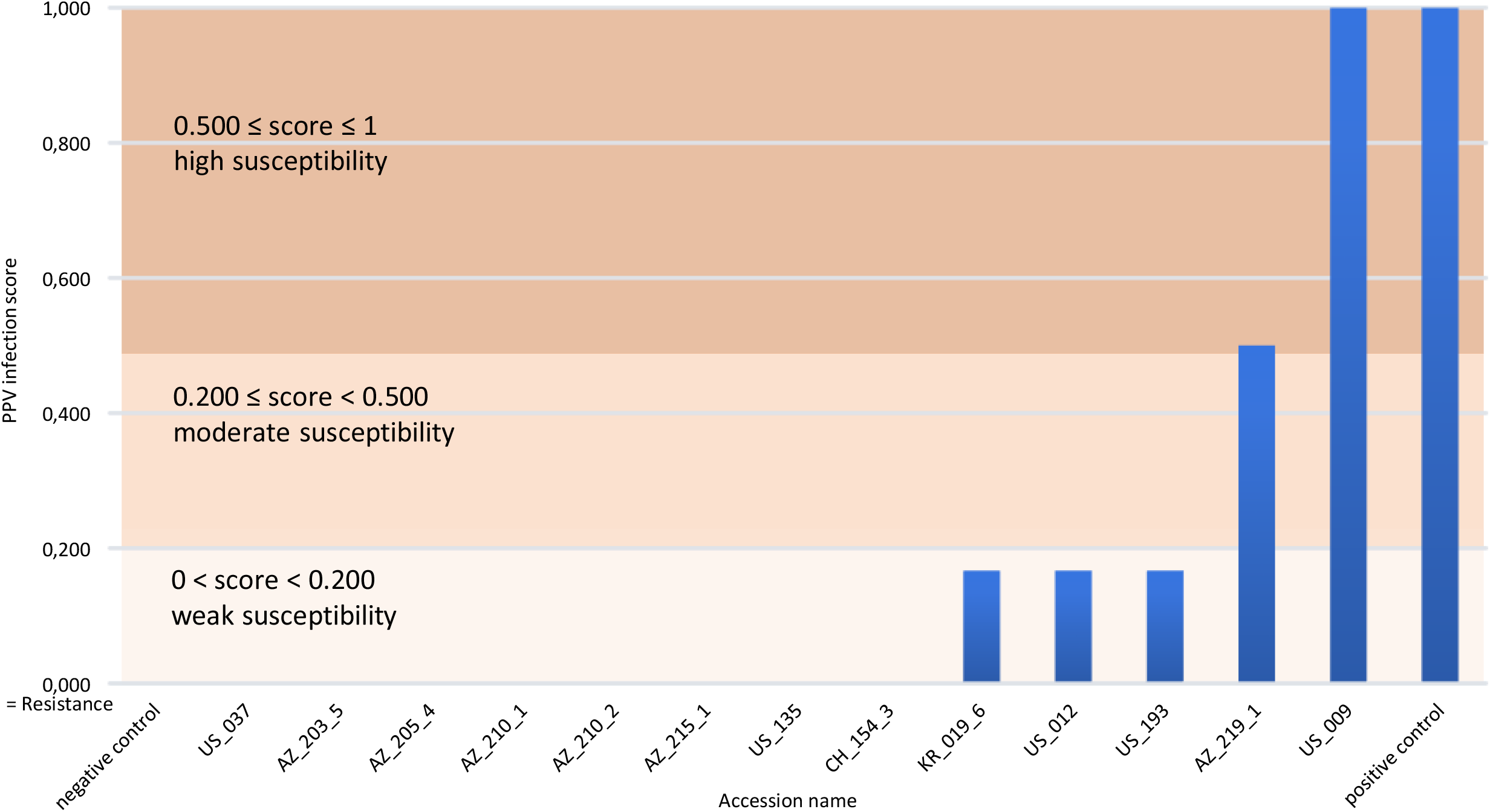
Average PPV infection score after successive cycles of phenotypic evaluation for the selected accessions (see Table 4). Resistance to PPV and the negative control are characterized by a null score (0). Three levels of susceptibility are categorized from a weak (0 < score < 0.200), then a moderate (0.200 ≤ score < 0.500) to a high susceptibility score (0.500 ≤ score ≤ 1). The positive control is set to 1.

## Discussion

Because of their small size (few kilobase pairs), viruses require (thus highjack) host proteins to complete their entire infectious cycle in the host plant, from viral RNA translation to virus movement, virion assembly and disassembly and viral replication (Garcia-Ruiz, 2018). In the case of PPV, two previous studies demonstrated the central role of components from the translation initiation complex, eIFiso4F, in the PPV compatible infection of stone fruit species (*Prunus sp*.) (Wang et al, 2012; Rubio et al, 2019). In the current study, the genetic diversity in the coding sequences of the recessive resistance gene *eIFiso4E* and of its counterpart, *eIF4E*, was analyzed in three *Prunus* crop species (apricots, almonds and peaches) and their wild related species from eleven geographic regions. The aim was to infer the evolutionary pattern of both genes involved in cellular translation initiation, to identify natural allele variations from susceptibility alleles and to evaluate the potential impact of these variations on *Prunus*/PPV interactions.

Such investigation of genetic diversity in natural populations for one or several genes is also called EcoTILLING. It is derived from the TILLING (Targeted Induced Local Lesion In Genome) strategy (Till et al., 2003) and was successfully used in the past as a non-transgenic reverse genetic approach in animals and plants to screen the natural diversity of targeted genes in many species (Barkley et al., 2008) especially for annual or pluriannual crop species as described by recent studies; in tomato (Upadhyaya et al., 2017), rice (Raja et al., 2017), wheat (Irshad et al., 2019) and cotton (Zeng et al., 2016). In perennial plants, there is one recent study dealing with the characterization of the nucleotide diversity found in the *TFL1* gene (Terminal Flower 1) in cultivated grapevine (Fernandez et al., 2014). The EcoTILLING strategy was also used in plants to successfully identify novel alleles for candidate genes involved in the resistance to diseases and specifically for the *eIF* susceptibility genes in melon (Nieto et al., 2007), pepper (Ibiza et al., 2010) and barley (Hofinger et al., 2011). Therefore, screening the genetic diversity of two *eIF* genes in *Prunus* species including both the crop and the non-domesticated germplasms is an original approach for perennial trees. It successfully identified novel alleles for both *eIF4E* and *eIFiso4E* genes in several *Prunus* species that could be associated with the resistance to PPV.

Polymorphism at both genes showed that the genetic diversity was globally higher among the wild related species than apricot, almond and peach crop species. Despite nucleotide variability present in the 1,397 accessions screened here, none of them displayed null alleles due to frameshift variations causing truncated proteins. Low proportions of non-synonymous variations (*dn/ds*<1 in the three main groups) likely indicated the removal of mutations that could be potentially deleterious or that could impair the proper functioning of the translation initiation complex. Likewise, Tajima’s *D* statistics for the both loci showed negative values, thus demonstrating an absence of neutrality between the mean pairwise difference and the number of segregating sites, explained by an abundance of low frequency haplotypes and the *dn/ds* ratios. Interestingly, the *eIFiso4E* coding sequence appeared more constrained than *eIF4E* with a *dn/ds* ratio consistently lower for *eIFiso4E* than for *eIF4E*, except for the wild undomesticated apricots. That would indicate a higher cost of non-synonymous mutations at the *eIFiso4E* locus than *eIF4E*. This result is consistent with the hypothesis that non-functional or silenced eIFiso4E alleles are lethal or deleterious in diploid *Prunus* growth (Rubio et al, 2019). Fitness cost of loss-of-function in eIFiso4E has been previously suspected for melon (Rodríguez-Hernández et al, 2012). The situation is different in wild apricots because of their self-incompatibility and thus an expected higher rate of heterozygosity would allow the occurrence of a higher number of non-synonymous mutations at the eIFiso4E locus.

Regarding the number of individuals with amino acid variations compared to those from the susceptible individuals, the estimators for: the haplotypic richness, the level of heterozygosity, the haplotype diversity (*Hd*) and the nucleotide diversity (*π*) were all significant strong indicators that *Prunus* related species actually constitute a reservoir of genetic diversity for alleles of interest. Three recent studies confirmed this view. In 2019, two consecutive studies showed that nucleotide diversity in peach wild related species was twice as much as that of peach landraces or peach improved cultivars (Velasco et al., 2016, Cao et al., 2019). More recently in 2021, Groppi et al. showed that undomesticated (wild) apricots from central Asia had global nucleotide diversity 1.5 times higher than cultivated European domesticated apricots.

When comparing genetic diversity within the *Prunoideae* taxonomic sections, we observed that eIF4E genetic diversity was the highest for the Amygdalus group while that of the eIFiso4E locus was highest for the Armeniaca group. In contrast for these two loci, members of the Persica group displayed the lowest value of genetic diversity (*π*), the lowest frequency of amino acid variations and the lowest level of heterozygosity. Consequently, the identification of potential variations in candidate genes is more likely to be successful firstly in related species and secondly in Amygdalus and Armeniaca than in Persica. Those accessions we found exhibiting allelic variations that potentially affected the overall protein conformation or the plant-potyvirus interaction domain(s) were all tested for susceptibility to sharka and eight out of thirteen were resistant to PPV, among which six were from the almond wild related species, *P. fenzliana* (5) and *P. tenella* (1), and one was from a peach related species, *P. davidiana*. Only one of them were from crop species (almond, *P. dulcis*)

Previous studies identified resistance to PPV in *P. davidiana*, in crop almond (*P. dulcis*) and in apricot (*P. armeniaca*) through quantitative trait phenotyping (Pascal et al, 2002; Hurtado et al., 2002, Vilanova et al., 2003; Decroocq et al., 2005, Marandel et al., 2009). One major locus controlling resistance to sharka in apricot has been mapped on linkage group 1 and named PPVres (Zuriaga et al., 2013, Decroocq et al., 2014, Mariette et al, 2016). While the *eIFiso4E* gene maps on the upper end of the chromosome 1 (Decroocq et al, 2005), it does not co-localize with PPVres. Therefore, finding eIFiso4E allelic variations impaired in *Prunus*-PPV interactions would provide new sources of resistance to sharka that could be combined with previously identified ones.

Provided that the observed resistance is indeed genetically controlled by the corresponding variations in the *eIFiso4E* coding sequence, our data provide new PPV resistant genitors. Regarding the amino acid variations; A28-and S157N are present in both susceptible (AZ_219_1, US_012, US_193) and resistant (AZ_203_5, AZ205_4, AZ210_1, AZ_210_2, AZ_215_1, US_037) individuals, thus, they are not correlated with the observed resistance phenotype. Nevertheless, further analyzes need to be performed for the amino acid variations V10-(US_135), A31P (AZ_205_4, AZ_210_1, AZ_215_1), D96E (US_135) and E187D (CH_154_3), all associated with PPV resistant individuals. Co-segregation between the above non-synonymous variations and response to PPV infection will have to be tested in F2 progenies because of the recessive nature of the resistance trait. However, this study provides initial insights on functional, genetic diversity and potential new sources of resistance to sharka.

## Statements

Appropriate permissions from responsible authorities for collecting and using *Prunus* samples from Central Asia, Caucasia and China were obtained by the local collaborators. The rest of the samples were kindly provided, with due authorizations, by the curators of the French INRAE Genetic Resources Centre (GRC), the US ARS-USDA repository, the Lednice germplasm collection; further details are available on their respective databases.

## Data availability

All the raw sequencing data generated during the current study were deposited in the SRA under project number: SUB12515842 and BioProject ID: PRJNA918999.

Supplemental tables are accessible to https://entrepot.recherche.data.gouv.fr/dataset.xhtml?persistentId=doi:10.57745/EYPRSU

## Acknowledgements

The authors wish to pay their respects to the late Dr Raul Karychev, former scientist of the Kazakh Research Institute of Horticulture and Viticulture (Almaty) who helped tremendously to initiate this work and did not have the chance to see its completion, even though he was an important contributor to this project.

We acknowledge the European FP7 IRSES-246795 “STONE”, the ANR-13-KBBE-0006 “COBRA” and the PRIMA FREECLIMB (# 1813-2) grants. S.L. was recipient of a Chinese Scholarship Council PhD grant. V.D. and S.L. thank the French Embassy in Beijing for accommodation and research support in China (Xu Guangqi program 2016-2018) and the Research Federation on Integrative Biology and Ecology of Bordeaux University for a travel grant.

The authors wish to acknowledge all the people who helped in collecting the samples. We thank the curators of the French Genetic Resources Centre (Marine Delmas), of the US ARS-USDA repository (John Preece) and of the Czech Horticultural repository of Lednice. We acknowledge the good and efficient care of the plants at the UMR BFP (INRAE) by Jean-Philippe Eyquard and Pascal Briard.

We also thank Drs Tatiana Giraud (ESE, CNRS-Univ. Paris Saclay) and Albert G. Abbott (University of Kentucky) for their critical review and proofreading of this manuscript.

**Supplemental tables available at https://entrepot.recherche.data.gouv.fr/dataset.xhtml?persistentId=doi:10.57745/EYPRSU**

**Table S1**: List of the individuals used in this study regarding the taxonomic classification in the *Prunus* genus, the geographic origin of the sampling, the subgroup classification used in this study and the eiF4E and eIFiso4E haplotypes obtained after the sequencing of the coding regions. Two haplotypes for one gene indicates the individual is heterozygous.

**Table S2**: Overall data from eIF4E gene for the three main groups (Armeniaca, Amygdalus and Persica) and their sub-groups respectively (crop species, wild forms and related species) concerning the sample size, the number of haplotypes and, in both coding and amino acid sequences the number of heterozygous individuals, the number of individuals with variation(s) and the haplotypic richness (allelic diversity) in **Table S2-A**; statistical analyses χ tests) with their associated *p-*values and classifications from the eiFiso4E gene between the three main groups (Armeniaca, Amygdalus and Persica) and the sub-groups (crop species, wild forms and related species) of each main group concerning the number of heterozygous individuals, the number of individuals with variation(s) and the haplotypic richness (allelic diversity) in both coding and amino acid sequences in **Table S2-B**. A *p*-value< 0.05 was considered significant and highlighted in grey with the following significance level: * for a *p-*value < 0.05, ** < 0.01 and *** < 0.0001 (highly significant).

**Table S3**: Overall data from eIFiso4E gene for the three main groups (Armeniaca, Amygdalus and Persica) and their sub-groups respectively (crop species, wild forms and related species) concerning the sample size, the number of haplotypes and, in both coding and amino acid sequences the number of heterozygous individuals, the number of individuals with variation(s) and the haplotypic richness (allelic diversity) in **Table S3-A**; statistical analyses (χ ^2^ tests) with their associated *p-*values and classifications from the eiFiso4E gene between the three main groups (Armeniaca, Amygdalus and Persica) and the sub-groups (crop species, wild forms and related species) of each main group concerning the number of heterozygous individuals, the number of individuals with variation(s) and the haplotypic richness (allelic diversity) in both coding and amino acid sequences in **Table S3-B**. A *p*-value < 0.05 was considered significant and highlighted in grey with the following significance level: * for a *p-*value < 0.05, ** < 0.01 and *** < 0.0001 (highly significant).

**Table S4-A**: Positions of amino acid variations along the eIFiso4E amino acid sequence (216 aa long) found in the Armeniaca group. Haplotypes derived from the amino acid reference sequence are indicated in the left column with the prefix _Arm (Armeniaca) followed by _P (Protein) with an incremented number from _P00 corresponding to the haplotype name given to the sequence of reference. Numbers of haplotypes regarding the main group, its sub-groups and the geographical area of origins of the individuals are indicated in the middle and the right side of the table respectively. Numbers in brackets refer to the total number of haplotypes in the group, each sub-group and according to the geographical area of the individuals. Numbers in square brackets indicate the frequency of the haplotype for the reference and/or the mostly frequent ones. The haplotype name, the amino acids and the number of haplotypes linked to the reference sequence are highlighted in grey. Positions of amino acid variations in the first interaction domain of the protein with the virus are indicated in blue. Haplotypes with an occurrence < 5% and those ones found homozygous are highlighted in pink and orange respectively. The symbol “-” indicate an amino acid deletion and “#” refers to a haplotype phenotyped after PPV infection in the greenhouse (see Table 4).

**Table S4-B**: Positions of amino acid variations along the eIFiso4E amino acid sequence (216 aa long) found in the Amygdalus group. Haplotypes derived from the amino acid reference sequence are indicated in the left column with the prefix _Amy (Amygdalus) followed by _P (Protein) with an incremented number from _P00 corresponding to the haplotype name given to the sequence of reference. Numbers of haplotypes regarding the main group, its sub-groups and the geographical area of origins of the individuals are indicated in the middle and the right side of the table respectively. Numbers in brackets refer to the total number of haplotypes in the group, each sub-group and according to the geographical area of the individuals. Numbers in square brackets indicate the frequency of the haplotype for the reference and/or the mostly frequent ones. The haplotype name, the amino acids and the number of haplotypes linked to the reference sequence are highlighted in grey. Positions of amino acid variations in the first interaction domain of the protein with the virus are indicated in blue. Haplotypes with an occurrence < 5% and those ones found homozygous are highlighted in pink and orange respectively. The symbol “-” indicate an amino acid deletion and “#” refers to a haplotype phenotyped after PPV infection in the greenhouse (see Table 4).

**Table S4-C:** Positions of amino acid variations along the eIFiso4E amino acid sequence (216 aa long) found in the Persica group. Haplotypes derived from the amino acid reference sequence are indicated in the left column with the prefix _Per (Persica) followed by _P (Protein) with an incremented number from _P00 corresponding to the haplotype name given to the sequence of reference. Numbers of haplotypes regarding the main group, its sub-groups and the geographical area of origins of the individuals are indicated in the middle and the right side of the table respectively. Numbers in brackets refer to the total number of haplotypes in the group, each sub-group and according to the geographical area of the individuals. Numbers in square brackets indicate the frequency of the haplotype for the reference and/or the mostly frequent ones. The haplotype name, the amino acids and the number of haplotypes linked to the reference sequence are highlighted in grey. Positions of amino acid variations in the first interaction domain of the protein with the virus are indicated in blue. Haplotypes with an occurrence < 5% and those ones found homozygous are highlighted in pink and orange respectively. The symbol “-” indicate an amino acid deletion and “#” refers to a haplotype phenotyped after PPV infection in the greenhouse (see Table 4).

**Table S5-A**: Positions of amino acid variations along the eIF4E amino acid sequence (234 aa long) found in the Armeniaca group. Haplotypes derived from the amino acid reference sequence are indicated in the left column with the prefix _Arm (Armeniaca) followed by _P (Protein) with an incremented number from _P00 corresponding to the haplotype name given to the sequence of reference. Numbers of haplotypes regarding the main group, its sub-groups and the geographical area of origins of the individuals are indicated in the middle and the right side of the table respectively. Numbers in brackets refer to the total number of haplotypes in the group, each sub-group and according to the geographical area of the individuals. Numbers in square brackets indicate the frequency of the haplotype for the reference and/or the mostly frequent ones. The haplotype name, the amino acids and the number of haplotypes linked to the reference sequence are highlighted in grey. Positions of amino acid variations in the first interaction domain of the protein with the virus are indicated in blue. Haplotypes with an occurrence < 5% and those ones found homozygous are highlighted in pink and orange respectively.

**Table S5-B:** Positions of amino acid variations along the eIF4E amino acid sequence (234 aa long) found in the Amygdalus group. Haplotypes derived from the amino acid reference sequence are indicated in the left column with the prefix _Amy (Amygdalus) followed by _P (Protein) with an incremented number from _P00 corresponding to the haplotype name given to the sequence of reference. Numbers of haplotypes regarding the main group, its sub-groups and the geographical area of origins of the individuals are indicated in the middle and the right side of the table respectively. Numbers in brackets refer to the total number of haplotypes in the group, each sub-group and according to the geographical area of the individuals. Numbers in square brackets indicate the frequency of the haplotype for the reference and/or the mostly frequent ones. The haplotype name, the amino acids and the number of haplotypes linked to the reference sequence are highlighted in grey. Positions of amino acid variations in the first interaction domain of the protein with the virus are indicated in blue. Haplotypes with an occurrence < 5% and those ones found homozygous are highlighted in pink and orange respectively.

**Table S5-C**: Positions of amino acid variations along the eIF4E amino acid sequence (234 aa long) found in the Persica group. Haplotypes derived from the amino acid reference sequence are indicated in the left column with the prefix _Per (Persica) followed by _P (Protein) with an incremented number from _P00 corresponding to the haplotype name given to the sequence of reference. Numbers of haplotypes regarding the main group, its sub-groups and the geographical area of origins of the individuals are indicated in the middle and the right side of the table respectively. Numbers in brackets refer to the total number of haplotypes in the group, each sub-group and according to the geographical area of the individuals. Numbers in square brackets indicate the frequency of the haplotype for the reference and/or the mostly frequent ones. The haplotype name, the amino acids and the number of haplotypes linked to the reference sequence are highlighted in grey. Positions of amino acid variations in the first interaction domain of the protein with the virus are indicated in blue. Haplotypes with an occurrence < 5% and those ones found homozygous are highlighted in pink and orange respectively.

**Table S6-A:** Predicted effects on protein function following the amino acid variations found on the eIFiso4E amino acid sequences in the Armeniaca main group. Predictions were processed online by three different softwares: 1-Meta-SNP prediction. The software gives a prediction score from 0 to 1, with a score > 0.5 the mutation is predicted “Disease” with an effect on protein function and/or conformation. From 0 to 0.5, the mutation is “Neutral”. “Nali” refers to the Number of Aligned sequences in the mutated site; 2-MutPred2 prediction. The software gives a prediction score from 0 to 1, with a score > 0.5 the mutation is predicted to affect protein function and/or conformation. In this case, indication about molecular mechanisms possibly affected with a *p*-value ≤ 0.05 and the associated probability found in the database are given; 3-PredictSNP. The software gives a confidence percentage that represents the percentage of expected accuracy if the amino acid variation is either neutral or deleterious on the protein function and/or conformation. All the predictor softwares are not able to calculate the potential effect of a deletion. An amino acid variation that potentially affects protein function with its prediction score is highlighted in grey. Position of an amino acid variation in the interaction domain I (DI) of the protein with the virus is indicated in blue. Haplotypes found with an occurrence < 5% in the main group was highlighted in pink. Haplotypes and individuals found homozygous are indicated in orange. The symbol “-” represents an amino acid deletion. The amino acid variation A127P in italics was included in the list of amino acid that possibly affect protein function because it had two scores very closed to the threshold and it was predicted to possibly affect the protein function in the Amygdalus group.

**Table S6-B:** Predicted effects on protein function following the amino acid variations found on the eIFiso4E amino acid sequences in the Amygdalus main group. Predictions were processed online by three different softwares: 1-Meta-SNP prediction. The software gives a prediction score from 0 to 1, with a score > 0.5 the mutation is predicted “Disease” with an effect on protein function and/or conformation. From 0 to 0.5, the mutation is “Neutral”. “Nali” referes to the Number of Aligned sequences in the mutated site; 2-MutPred2 prediction. The software gives a prediction score from 0 to 1, with a score > 0.5 the mutation is predicted to affect protein function and/or conformation. In this case, indication about molecular mechanisms possibly affected with a *p*-value ≤ 0.05 and the associated probability found in the database are given. 3-PredictSNP. The software gives a confidence percentage that represents the percentage of expected accuracy if amino acid variation is either neutral or deleterious on the protein function and/or conformation. All the predictor softwares are not able to calculate the potential effect of a deletion. An amino acid variation that potentially affects protein function with its prediction score is highlighted in grey. Positions of amino acid changes in the interaction domains I (DI) and II (DII) of the protein with the virus are indicated in blue and green respectively. Haplotypes found with an occurrence < 5% in the main group was highlighted in pink. Haplotypes and individuals found homozygous are indicated in orange. The symbol “-” represents an amino acid deletion. The amino acid variation K204I in italics was included in the list of amino acid that possibly affect protein function because it had two scores very closed to the threshold.

**Table S6-C**: Predicted effects on protein function following the amino acid variations found on the eIFiso4E amino acid sequences in the Persica main group. Predictions were processed online by three different softwares: 1-Meta-SNP prediction. The software gives a prediction score from 0 to 1, with a score > 0.5 the mutation is predicted “Disease” with an effect on protein function and/or conformation. From 0 to 0.5, the mutation is “Neutral”. “Nali” refers to the Number of Aligned sequences in the mutated site; 2-MutPred2 prediction. The software gives a prediction score from 0 to 1, with a score > 0.5 the mutation is predicted to affect protein function and/or conformation. In this case, indication about molecular mechanisms possibly affected with a *p*-value ≤ 0.05 and the associated probability found in the database are given; 3-PredictSNP. The software gives a confidence percentage that represents the percentage of expected accuracy if amino acid change is either neutral or deleterious on the protein function and/or conformation. All the predictor softwares are not able to calculate the potential effect of a deletion. An amino acid variation that potentially affects protein function with its prediction score is highlighted in grey. Position of amino acid variation in the interaction domain II (DII) of the protein with the virus is indicated in green. Haplotypes found with an occurrence < 5% in the main group is highlighted in pink. Haplotypes and individuals found homozygous are indicated in orange. The symbol “-” represents an amino acid deletion. The amino acid variation A127P in italics was included in the list of amino acid that possibly affect protein function because it had two scores very closed to the threshold and was predicted to affect the protein function in the Amygdalus group.

**Table S7:** Scoring of PPV infection after DAS-ELISA tests for the selected accessions in table 4. Score = 1 indicates a positive DAS-ELISA test, score = 0 indicates a negative DAS-ELISA test. Empty boxes indicate no measurement. For each accession, the score in the grey box was used to perform Figure 4.

